# Discovery of essential kinetoplastid-insect adhesion proteins and their function in *Leishmania*-sand fly interactions

**DOI:** 10.1101/2023.09.29.560188

**Authors:** Ryuji Yanase, Katerina Pružinová, Edward Rea, Flávia Moreira-Leite, Atsushi Taniguchi, Shigenori Nonaka, Jovana Sádlová, Barbora Vojtkova, Petr Volf, Jack D. Sunter

## Abstract

***Leishmania* species, members of the kinetoplastid parasites, cause leishmaniasis, a neglected tropical disease, in millions of people worldwide^1^. *Leishmania* has a complex life cycle with multiple developmental forms, as it cycles between a sand fly vector and a mammalian host; understanding their life cycle is critical to understanding disease spread^2^. One of the key life cycle stages is the haptomonad form, which is attached to the insect through its flagellum. This adhesion, which is conserved across kinetoplastid parasites, is implicated to have an important function within their life cycles and hence on disease transmission^3–5^. Here, we discovered kinetoplastid-insect adhesion proteins (KIAPs), which are localised in the attached haptomonad flagellum. Deletion of these KIAPs impaired cell adhesion *in vitro* and prevented *Leishmania* from colonising the stomodeal valve in the sand fly, without affecting cell growth. This result will provide important insights for a comprehensive understanding of the *Leishmania* life cycle.**

---

The specific interaction and adhesion of a pathogen to a tissue is a common strategy enabling immune evasion, maintenance of infection, and life cycle progression, such as sequestration of *Plasmodium falciparum* in the vasculature^6^. By anchoring themselves, pathogens can modify their environment, with *Yersinia pestis* and the *Leishmania* parasites secreting an extracellular matrix that partially blocks the gut of their vectors leading to increased feeding and hence transmission^3,7,8^.

The *Leishmania* haptomonad form is an important example of strong adhesion between a vector and pathogen^9^. *Leishmania* spp. are the causative agent of leishmaniasis^1^. These flagellate parasites are transmitted between mammalian hosts by sand flies and within the fly there are five developmental stages. Initial colonisation of the sand fly occurs through an infected blood meal after which amastigotes differentiate into motile promastigotes. These promastigotes migrate to the anterior midgut differentiating to human-infective metacyclics and haptomonads attached to the stomodeal valve^2^. Destruction of the stomodeal valve by *Leishmania* results in regurgitation of parasites into the mammalian host during the next bloodmeal^8,10^. Understanding the molecular cell biology of these cryptic interactions is key to understanding colonisation and transmission dynamics of these important pathogens.

Haptomonad adhesion occurs via the heavily modified parasite flagellum, in which an attachment plaque is positioned adjacent to the flagellum membrane from which filaments extend towards the cell body and specific connections to the stomodeal valve are formed^11^. *Leishmania* is closely related to other kinetoplastid parasites that cause disease in humans. Adhesion of these parasites through their flagellum to their insect vector is an important part of their life cycle^3,12^ and is associated with the generation of mammalian infective forms, especially for *Trypanosoma cruzi* and *T. congolense*^4,5^. Despite the importance of these attached forms the molecular makeup of them remains cryptic. Here, we identified the first proteins of the *Leishmania* attachment plaque and showed they are necessary for its assembly and the colonisation of the sand fly stomodeal valve. This opens up new avenues to dissect the biology of these critical forms for all kinetoplastid parasites.

To dissect the components of the *Leishmania* haptomonad attachment plaque, we used comparative proteomics to identify proteins enriched in the attached haptomonad flagellum in comparison to the unattached promastigote flagellum (Fig. 1a). We identified 371 proteins enriched in the attached haptomonad flagellum (Supplementary Dataset 1). To refine our candidate list, we used TrypTag^13^, the genome-wide protein localisation resource for the related parasite *Trypanosoma brucei* to remove proteins whose orthologs localised to organelles unlikely to be involved in adhesion (Supplementary Dataset 2). 20 candidate proteins were taken forward, which we endogenously tagged with mNeonGreen (mNG) and examined their localisation in promastigotes and haptomonads (Extended Data Fig. 1). We identified three proteins that were enriched at the enlarged tip of the attached flagellum and named these proteins Kinetoplastid-Insect Adhesion Proteins (KIAPs) 1–3 (Fig. 1b).

**Figure 1.**
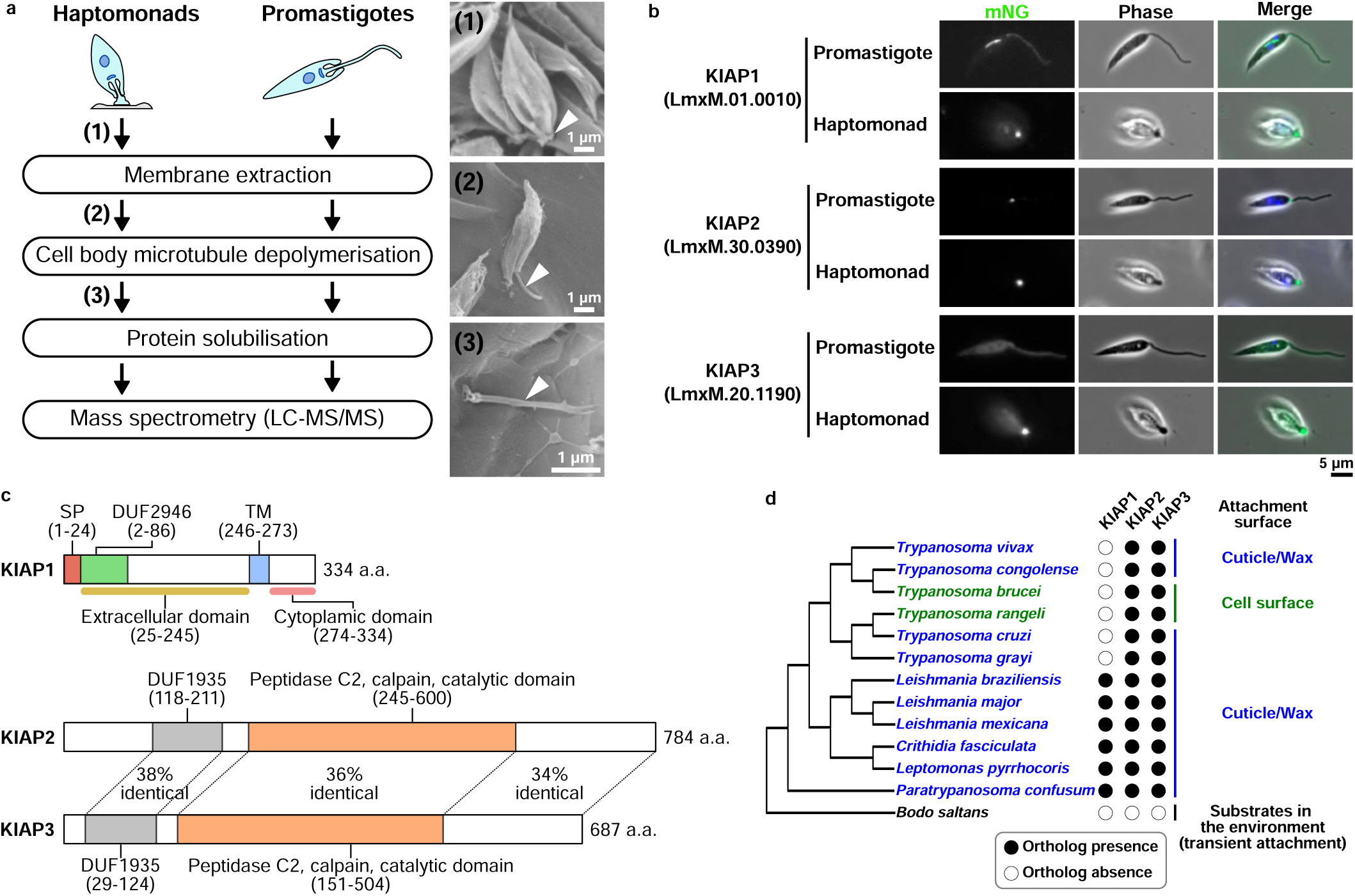
KIAPs identified through comparative proteomics. **a**, Experimental flow of sample preparation for the comparative proteomic using *in vitro* attached haptomonads and non-attached promastigotes. Scanning electron microscopy images show an intact *in vitro* haptomonad (1), a haptomonad after membrane extraction (2) and an attached flagellum remained on the substrate after cell body microtubule depolymerisation (3; the numbers correspond to the ones in the experimental flow, and each sample was prepared in the step indicated by the corresponding number). Arrowheads: attached flagellum. **b**, Localisation of mNeonGreen (mNG) tagged KIAPs in promastigote and *in vitro* haptomonad cells. **c**, Domain structures of *L. mexicana* KIAPs. Domain names and ranges in the amino acid sequence are shown above coloured boxes. The amino acid (a.a.) length of each protein is indicated to the right of each structure. SP: signal peptide, DUF: domain of unknown function, TM: transmembrane domain. **d**, Phylogenetic tree of kinetoplastids based on 18S rRNA sequences showing conservation of KIAPs across the kinetoplastids. The presence (filled circle) or absence (open circle) of an ortholog of a gene family are indicated on the right-hand side of the tree. The species indicated in blue are known to adhere to cuticle or wax layers in the insect vector and the species indicated in green are known to adhere to the cell membrane in the insect vector. *Bodo saltans* exhibits transient adhesion to substrates in the environment. Branch lengths do not represent evolutionary time.

KIAP1 is a predicted type 1 membrane protein, with a signal peptide and transmembrane domain positioned towards the C-terminus (Fig.1c). The N-terminus of KIAP1 contains a DUF2946 domain and the AlphaFold structural prediction of the extracellular region is similar to the C-type lectin domain of tetranectin (Extended Data Fig. 2). This suggests that a carbohydrate moiety likely plays a partial role in mediating *Leishmania* adhesion. The origin of this carbohydrate is unclear; within the sand fly, the cuticle contains N-acetyl-glucosamine but this is potentially inaccessible below its wax surface, while in culture the serum may provide carbohydrate for adhesion. *Leishmania* also secretes proteophosphoglycans^14^ and these could act as the substrate for KIAP1 to bind. A similar strategy is used by fungal plants and insects pathogens, where they secrete substrates which they use to anchor themselves to the cuticle^15^. However, *in vitro Leishmania* adheres to hydrophobic substrates and may form adhesion through hydrophobic interactions on the waxy surface of the stomodeal valve, as suggested for *T. cruzi* adhesion^16^.

KIAP2 and KIAP3 have a similar domain structure, with an N-terminal DUF1935 domain and a central calpain C2 domain (Fig. 1c). The calpain domain is catalytically inactive as it lacks the critical cysteine residue and is found in many other important cytoskeletal proteins in the trypanosomatids^17–19^.

A reciprocal best BLAST analysis revealed that KIAP1 is present in all *Leishmania* species and other species including *Crithidia* and the early branching *Paratrypanosoma* but not present in *Trypanosoma* spp. and *Bodo saltans*, a free-living relative of the kinetoplastid parasite, which transiently adheres to substrates through its flagellum (Fig. 1d). KIAP2 and KIAP3 were conserved across the kinetoplastids except in *Bodo saltans* (Fig. 1d). This shows that the KIAPs are well conserved across the kinetoplastids, and likely represent a conserved set of components for parasite adhesion to their vectors.

*Leishmania* adhesion proceeds through a series of defined steps, from initial adhesion, to flagellum disassembly and final maturation of the plaque^11^. To understand the point at which the different KIAPs appeared, we examined cells at different stages of adhesion (Fig. 2a–c). KIAP1::mNG in a promastigote cell localised to the lysosome with a weak signal along the flagellum membrane, whereas mNG::KIAP3 had a weak signal throughout the cytoplasm and the flagellum. During adhesion both proteins behaved similarly, with bright spots of KIAP1::mNG and mNG::KIAP3 observed at points along the flagellum, coinciding with membrane deformations (arrowheads; Fig. 2a, c). As the flagellum disassembled, KIAP1::mNG and mNG::KIAP3 became concentrated in the enlarged region of the flagellum adjacent to the anterior cell tip (Fig. 2a, c). In contrast, in the promastigote, KIAP2::mNG localised to a small point within the flagellum, as the flagellum emerges from the cell body (Fig. 2b). During adhesion, cells were seen in which the KIAP2::mNG signal extends along the flagellum for a short distance or an additional focus of KIAP2::mNG was observed associated with a membrane deformation (arrowhead; Fig. 2b). As the flagellum dissembled, the KIAP2::mNG signal strength increased (Fig. 2b). These results demonstrate that KIAP1, KIAP2 and KIAP3 appear at the earliest stages of parasite adhesion and likely have a role in the initial interaction.

**Figure 2.**
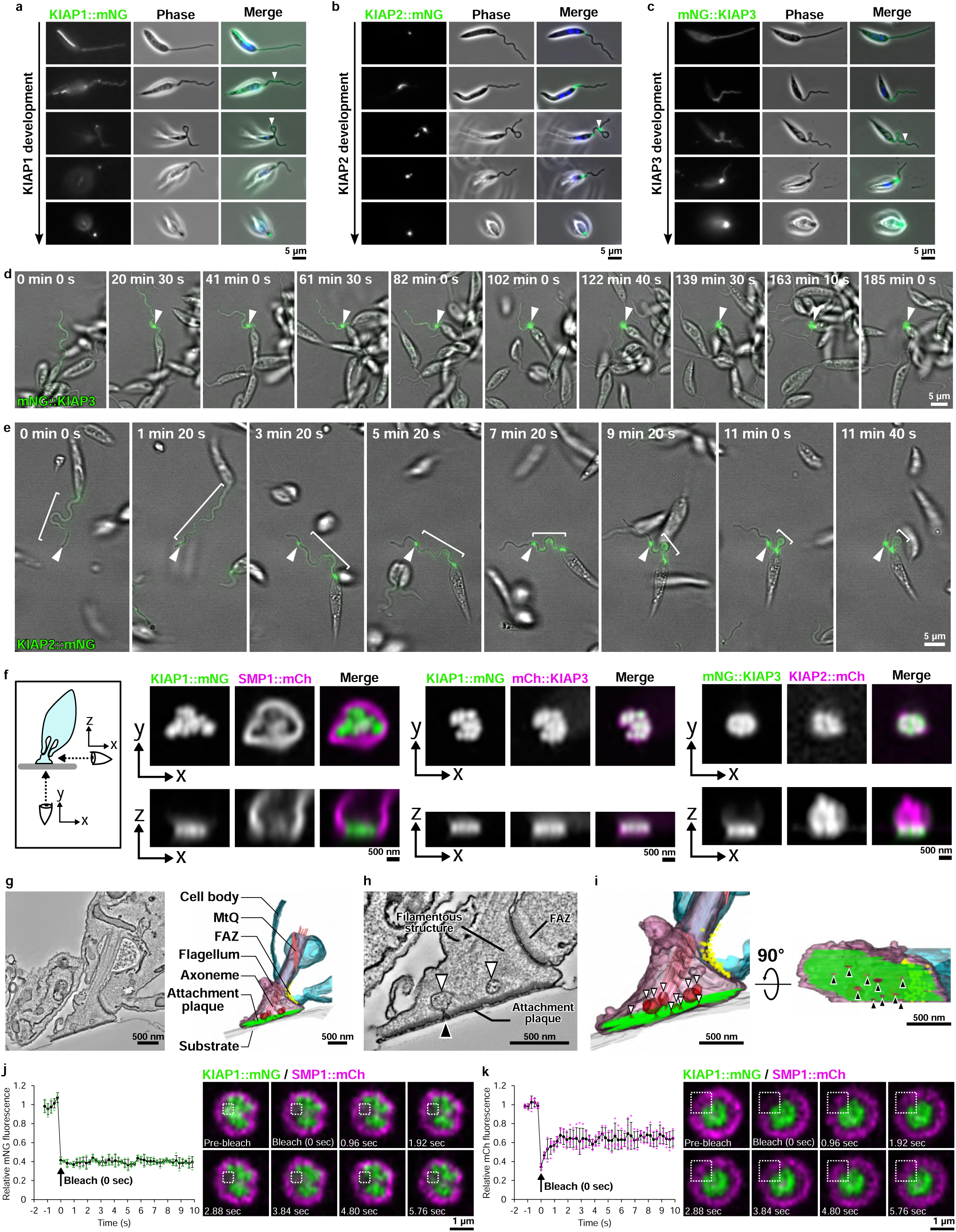
KIAPs have distinct localisation and development pattern during haptomonad adhesion. **a–c**, Development of mNG-tagged KIAP1 (a), KIAP2 (b) and KIAP3 (c) during differentiation from a promastigote cell to a haptomonad cell *in vitro*. Arrowheads: bright spots of KIAP1::mNG (a) or mNG::KIAP3 (c), or an additional focus of KIAP2::mNG (b) associated with a deformation of the flagellum membrane. **d**, Sequential frames (at ∼20 min intervals) from a time-lapse video of adhesion of a haptomonad cell and development of KIAP3 N-terminally tagged with mNG. Note that the starting point of adhesion (arrowheads) remains fixed until the completion of adhesion. **e**, Sequential frames (at ∼2 min intervals) from a time-lapse video of adhesion of a haptomonad cell and development of KIAP2 C-terminally tagged with mNG. The ranges indicated by the white bars show the range of KIAP2::mNG signal in the flagellum. Note that the starting point of adhesion (arrowheads) remains fixed during the adhesion process. **f**, Z-stack confocal microscopy analysis of KIAP localisation in an attached flagellum. The schematic diagram shows viewing directions of an attached flagellum (bottom view: X–Y, side view: X–Z). The images on the left shows the localisation of KIAP1::mNG (green) and the flagellar membrane marker, SMP1::mCherry (mCh; magenta). The images on the middle shows the localisation of KIAP1::mNG (green) and mCh::KIAP3 (magenta). The images on the right shows the localisation of mNG::KIAP3 (green) and KIAP2::mCh (magenta). The viewing directions are indicated in the bottom left-hand corner of the images. **g**, A slice of a serial tomogram and 3D reconstruction of an *in vitro* haptomonad cell. Vesicles near the attachment plaque (green) are shown in red. The name of each structure is given in the 3D reconstruction. MtQ: microtubule quartet. FAZ: flagellum attachment zone. **h**, Magnified view of the tomographic slice showing that multiple vesicles (white arrowheads) are seen near the attachment plaque and the plaque is interrupted at the fusion of the vesicle and flagellar membrane (black arrowhead). **i**, Magnified view of the 3D reconstruction showing multiple vesicles (white arrowheads) are seen near the attachment plaque (green) and the plaque is interrupted at the fusion of the vesicle and flagellar membrane (black arrowheads). Side (left) and bottom (right) views of the attached flagellum are shown. **j**, **k**, FRAP experiments of KIAP1::mNG (j) and SMP1::mCh (k). The relative fluorescence intensity changes before and after photobleaching are shown in the line graphs, with the average fluorescence intensity before photobleaching as 1. Data represent mean ± SD (n = 4 independent experiments). Values from each experiment are shown with green (j) or magenta (k) dots, respectively. The timing of photobleaching is indicated by an arrow in the graph. Sequential frames (at ∼1 sec intervals) of confocal microscopy observation of KIAP1::mNG and SMP1::mCh are shown. The area of photobleaching is indicated with dotted white boxes.

Next, we used time lapse analysis to follow changes in KIAPs localisation during adhesion (Fig. 2d, e). We followed cells expressing the individual KIAPs tagged with mNG and additionally generated a cell line expressing KIAP1 endogenously tagged with mNG and KIAP3 tagged with mCherry (mCh). For KIAP1 and KIAP3, the first indication of adhesion was the localisation of these proteins to a region of flagellum membrane deformation (Fig. 2d, Extended Data Fig. 3a and Supplementary Video 1 and 2). Over time, KIAP1 and 3 signal intensity increased at this region, with the flagellum disassembling until the mature haptomonad is formed. When the initial adhesion point occurred distal to the anterior cell tip both KIAP1 and KIAP3 were associated with the flagellum membrane deformation. This adhesion point remained fixed to the substrate, whilst the flagellum moved freely resulting in the movement of the cell body to this adhesion point (Fig. 2d, Extended Data Fig. 3a and Supplementary Video 1, 2).

During adhesion, KIAP2::mNG was observed to localise along the flagellum connecting the KIAP2 focus in the flagellum at the anterior cell tip to a second KIAP2 focus that coincided with a point of membrane deformation (Fig. 2e and Supplementary Video 3). The KIAP2::mNG focus associated with the adhesion point remained fixed to the surface and the cell body moved towards this point, with a concomitant reduction in the KIAP2 signal length between two foci (Fig. 2e and Supplementary Video 3). Yet, if adhesion occurred as the flagellum emerges from the cell body, KIAP2 localised only to this position (Extended Data Fig. 3b and Supplementary Video 4). Remarkably, this reveals that during adhesion the cell body can move towards the initial adhesion point, potentially requiring a traction force. Moreover, there appears to be two populations of KIAP2, one localised to the flagellum as it emerges from the cell body and another mobile within the flagellum.

To investigate the position of the KIAPs within the attached flagellum in more detail, we used super-resolution confocal microscopy. We generated cells expressing KIAP1 endogenously tagged with mNG and SMP1, a flagellum membrane marker, fused to mCh and examined the localisation of these proteins in attached cells (Fig. 2f). KIAP1 localised directly next to the surface on which the parasite was attached, forming a narrow band when viewed from the side and a patchy discontinuous surface when viewed from beneath. SMP1 was excluded from the attachment interface and this suggests that the membrane within the attachment plaque has a distinct composition to the rest of the flagellum membrane.

We then generated a set of cell lines with combinations of KIAPs endogenously tagged with either mNG or mCh to determine their relative positions (Fig. 2f). mCh::KIAP3 had a similar localisation to KIAP1::mNG, forming a band adjacent to the glass, with a patchy lateral distribution. While, KIAP2::mCh localised along the flagellum from adjacent to the plaque, as defined by KIAP3 localisation, to the cell body. This suggests that KIAP1 and 3 are found within the attachment plaque and KIAP2 within the filaments that run from the plaque towards the cell body (Fig. 2f).

Given the non-uniform distribution of KIAP1 and 3 across the attachment plaque, we revisited our electron tomograms of *Leishmania* attached in the sand fly and to plastic^11^ (Fig. 2g–i and Extended Data Fig. 3c–f). We were readily identified breaks within the attachment plaque in the tomograms, correlating with our light microscopy (Fig. 2h, i and Extended Data Fig. 3d, e). Across these five plaques, 23 of the 27 breaks we found were associated with an invagination of the flagellum membrane. These invaginations likely represent the fusion of vesicles within the flagellum that deliver components for the assembly of the plaque and/or material to be secreted into the local environment to influence parasite infection. Interestingly, the material within the vesicles in the sand fly haptomonads has a similar filamentous appearance to the extracellular material between the parasites (Extended Data Fig. 3d, f). This may represent the secretion of the filamentous proteophosphogylcan or acid phosphatase both known to be released by *Leishmania*^14^.

Our electron tomography shows that once assembled the attachment plaque is a highly organised structure, and despite the small breaks associated with membrane dynamics, we hypothesised that there would be limited movement and turnover of the KIAPs. We used fluorescence recovery after photobleaching to investigate the movement of KIAP1::mNG within the plaque (Fig. 2j and Extended Data Fig. 3g). After photobleaching of KIAP1::mNG, we did not observe any recovery of signal either over a short period or longer period, whereas the SMP1::mCh signal, which is restricted to the membrane outside of the plaque rapidly recovered (Fig. 2k). This shows, as predicted, that there is limited movement and turnover of KIAP1 within the attachment plaque and potentially reflects KIAP1 binding to its substrate. The capacity of haptomonads to differentiate is unknown but the stability of the attachment plaque suggests that the attached cell is terminally differentiated. Interestingly, we previously observed that ∼1% of haptomonads were dividing^11^, which may generate a free-swimming daughter and such asymmetric differentiation divisions are found in other kinetoplastid parasites^20^.

The highly structured nature of the attachment plaque is likely the result of an ordered assembly process. To investigate the dependency relationships between the different KIAPs, we generated cell lines in which either KIAP1, KIAP2 or KIAP3 were deleted and one of the other two were endogenously tagged with mNG (Fig. 3a–c). Successful null mutant generation was confirmed by PCR (Extended Data Fig. 4a, b). The loss of any of the KIAPs led to a limited number of cells adhering, and we examined KIAP localisation in these cells. The localisation of KIAP2::mNG appeared unaffected by the loss of either KIAP1 and KIAP3, with the protein localised at the point of flagellum emergence from the cell body, with occasional extensions along the flagellum seen during adhesion (white bars; Fig. 3a, c). The deletion of either KIAP2 and KIAP3 affected the localisation of KIAP1::mNG during adhesion. Without KIAP2 and KIAP3, KIAP1::mNG did not become concentrated in a focus within the flagellum but was instead distributed throughout the flagellum and cell body, and in cells with a short flagellum there was no flagellum signal (Fig. 3b, c). In KIAP1 and KIAP2 null mutants, mNG::KIAP3 did form bright spots in the flagellum (arrowheads; Fig. 3a, b), as seen in the parental cells; however, in cells with a shorter flagellum there was little flagellum signal (Fig. 3a, b). This suggests that KIAP1 localisation is dependent on KIAP2 and KIAP3, whereas KIAP3 is initially able to localise to points of adhesion but this localisation was not stable without KIAP1 and KIAP2.

**Figure 3.**
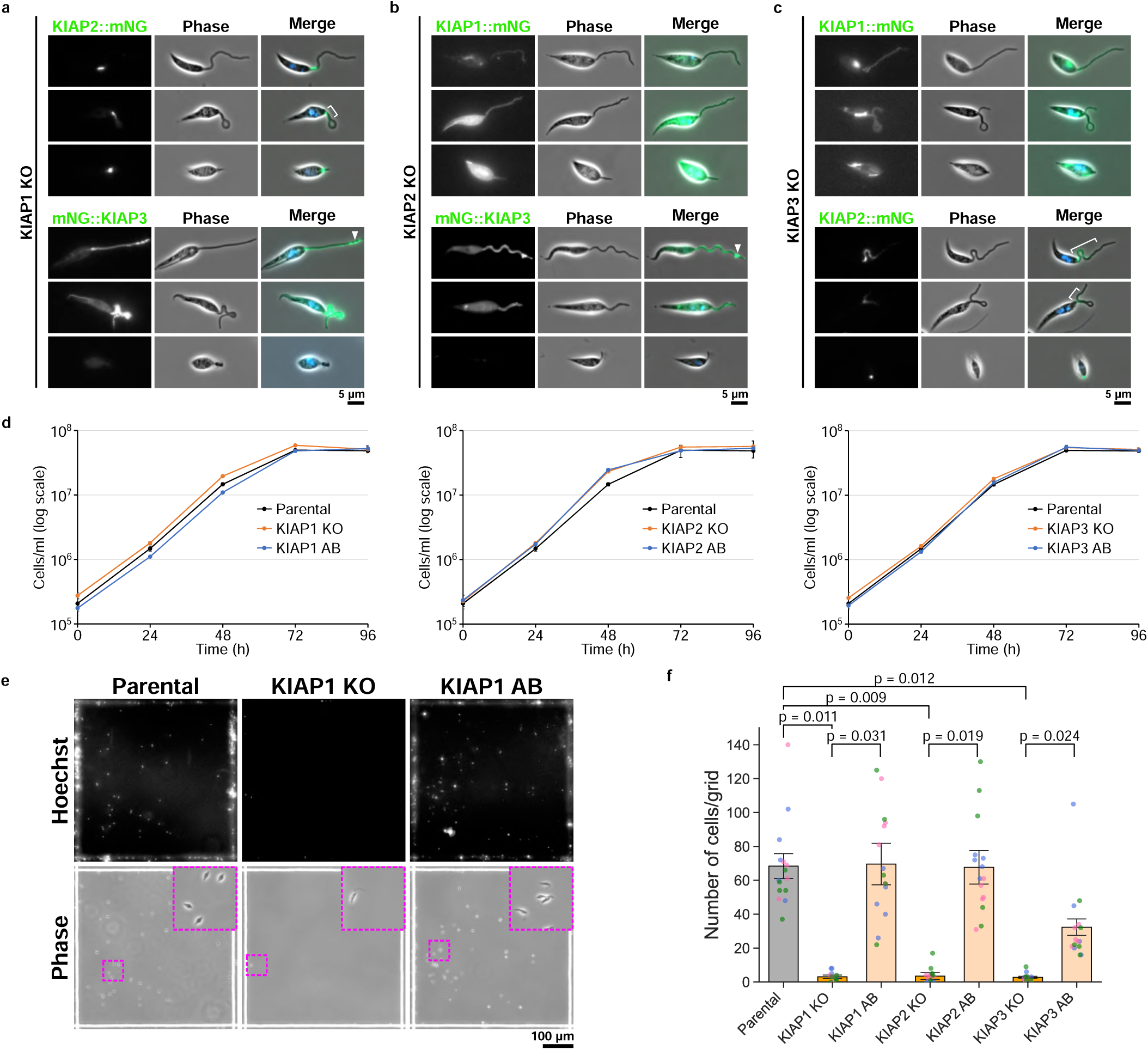
Deletion of KIAPs reduced parasite adhesion *in vitro*. **a–c**, Localisation of mNG-tagged KIAPs in KIAP1 (a), KIAP2 (b) or KIAP3 (c) KO *L. mexicana* cell lines. White bars: extensions of mNG::KIAP2 along the flagellum. Arrowheads: bright spots of mNG::KIAP3. **d**, Growth curves of parental and KIAP1 (left), KIAP2 (middle) and KIAP3 (right) gene knockout (KO) and add back (AB) cell lines. Data represent mean ± SD (n = 3 independent experiments). **e**, Hoechst fluorescence and phase images of parental, KIAP1 KO and AB *Leishmania* cells induced adhesion for 24 h on a gridded glass coverslip. Top; Hoechst images. Bottom; phase images. Insets show a magnified view of the magenta dotted square area. **f**, Quantification of the number of attached cells per grid area for parental and KIAPs KO and AB *L. mexicana* cell lines. Data represent mean ± SEM (n = 3 independent experiments). The blue, pink and green dots represent measurement from three independent experiments, respectively. P-values calculated using two-tailed Welch’s t-test, which are shown above the graph.

KIAP loss appeared to cause a catastrophic failure to generate mature attached cells and we sought to quantify this effect. The growth of KIAP1, KIAP2 or KIAP3 null mutants as promastigotes was unaffected (Fig. 3d), but the loss of any of the KIAPs resulted in a dramatic reduction in adhesion to glass (Fig 3e, f). To confirm that the loss of adhesion was specific to KIAP deletion, we generated add back cells in which a tagged version of KIAP1, KIAP2, or KIAP3 was introduced into the respective null mutant, with their expression and localisation to the enlarged flagellum tip of the haptomonad confirmed by fluorescence microscopy (Extended Data Fig. 4c). There was an increase in the number of attached cells observed for the add back cell lines in comparison to the null mutants (Fig. 3e, f), confirming that KIAP1, KIAP2 and KIAP3 are necessary for adhesion *in vitro*.

To investigate the importance of KIAPs for the formation of haptomonads in the sand fly, we infected flies with the parental, null mutant and add back cell lines for KIAP1 and KIAP3, and examined the distribution of parasites on day 6 and 9 post-blood meal (PBM; Fig. 4). There were minimal differences in the infection rates and loads between the different cell lines and all developed heavy late-stage infections (Fig. 4a); however, large differences were found in the location of the parasites (Fig. 4b). While 70–90% of sand flies had the stomodeal valve colonised by the parental and add back cells by day 9 PBM, no colonisation of the valve was observed for the KIAP1 and KIAP3 null mutants. The null mutants migrated to the cardia region but did not attach to the stomodeal valve.

**Figure 4.**
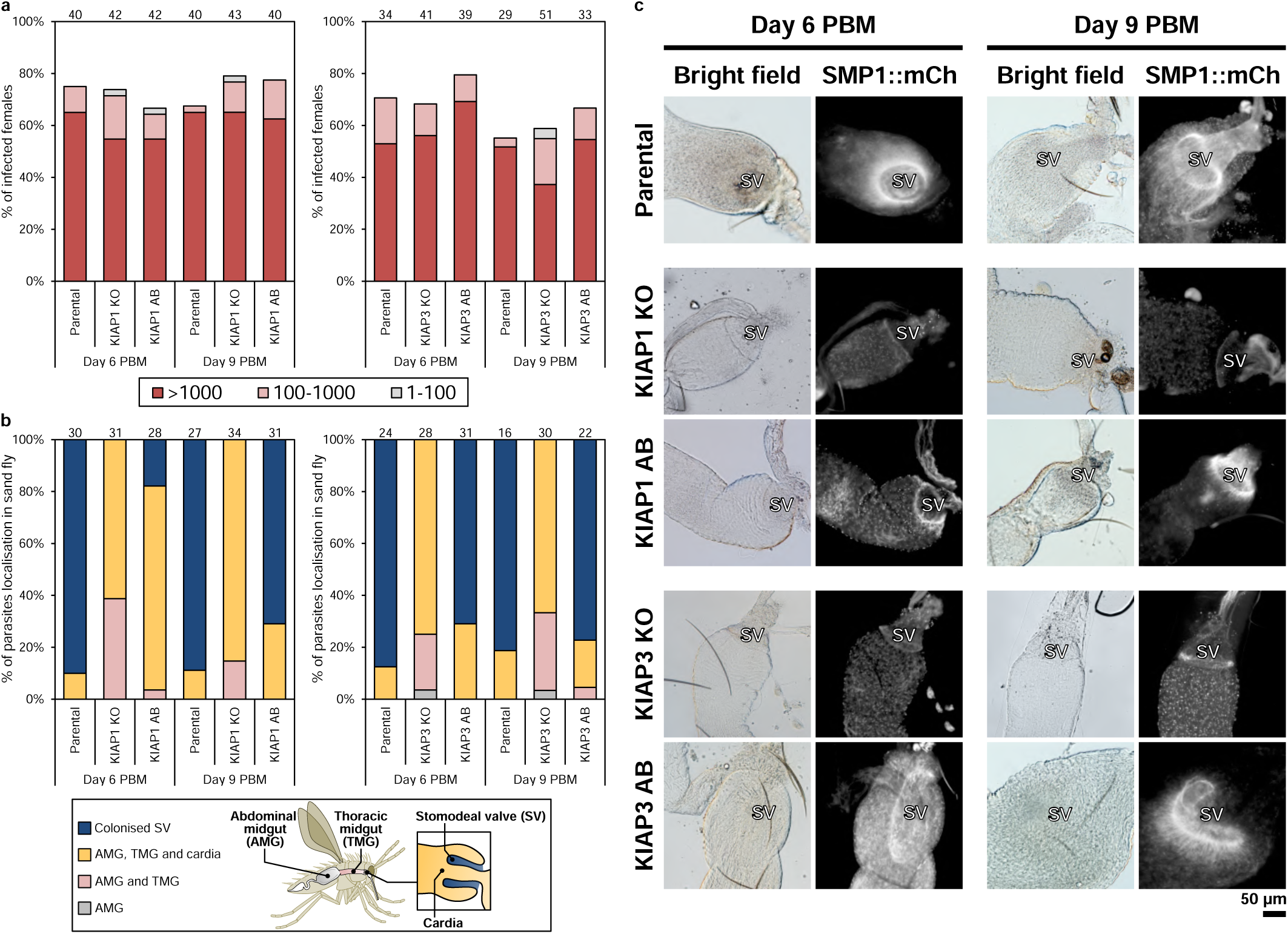
KIAP1 and KIAP3 were essential for stomodeal valve colonisation in the sand fly. **a**, Infection rates and intensities of infections of parental and KIAP1 (left) or KIAP3 (right) KO and AB *L. mexicana* cells on days 6 and 9 post blood meal (PBM). Numbers above the bars indicate the number of dissected female sand flies. Infection intensities were evaluated as light (<100 parasites/gut; grey), moderate (100–1000 parasites/gut; pink) and heavy (>1000 parasites/gut; red). The percentages obtained from two independent experiments are shown in the graphs. **b**, Localisation of infections of parental and KIAP1 (left) or KIAP3 (right) KO and AB *L. mexicana* cells on days 6 and 9 PBM. Numbers above the bars indicate the number of evaluated (positive) female sandflies. The schematic diagram shows the name and location of each part of a sand fly gut. The percentages obtained from two independent experiments are shown in the graphs. **c**, Bright field and fluorescence images of dissected sand fly guts on days 6 (left) or 9 (right) PBM where parental, or KIAP1 or KIAP3 KO and AB *L. mexicana* cells expressing SMP1::mCh infected. The location of the stomodeal valve is shown with SV.

We imaged the valves from infected sand flies (Fig. 4c). The parasites expressed the flagellum membrane marker SMP1 fused to mCherry, enabling us to determine their position. For both the parental and add back cells *Leishmania* parasites were associated with the stomodeal valve, whereas for the KIAP1 and KIAP3 null mutants there was a near complete loss of SMP1::mCh signal associated with the valve (Fig 4c). Overall, this shows that both KIAP1 and KIAP3 are necessary for colonisation of the stomodeal valve in the sand fly, correlating with the loss of adhesion *in vitro*.

In conclusion, we have identified and functionally characterised the first proteins necessary for the generation of the attachment plaque and adhesion of *Leishmania* to the stomodeal valve in the sand fly. The conservation of these proteins across the kinetoplastids suggests a common mechanism of adhesion and attachment plaque generation and provides an important insight into our understanding of the development and differentiation of these parasites in their insect vectors. Furthermore, since attachment is necessary for mammalian infection for a number of these parasites, our work is an important step to the development of transmission-blocking strategies.

## Methods

### Cell culture

Cas9T7 *L. mexicana* (WHO strain MNYC/BZ/1962/M379, expressing Cas9 and T7 RNA polymerase) promastigotes were grown at 28°C in M199 medium with 10% foetal calf serum, 40 mM HEPES-HCl (pH 7.4), 26 mM NaHCO_3_ and 5 µg/ml haemin. Cells were maintained in logarithmic growth by regular subculturing.

### *In vitro* haptomonad cell adhesion

Axenic haptomonad cells were generated by culturing 1 × 10^6^ cells/ml promastigotes on 13 mm round Thermanox plastic coverslips (Nalgene Nunc International, Rochester, NY) scratched with sandpaper and sterilised with 100% ethanol (for proteomics and electron microscopy) or gridded glass coverslips grid-500 (iBidi, Gräfelfing, Germany) which were cut into small pieces of ∼5 × 5 mm and sterilised with 100% ethanol (for widefield epifluorescence microscopy) in a 24 well plate with 1 ml of M199 medium at 28°C with 5% CO_2_ for 72 h with M199 medium being replaced every 24 h (for proteomics and electron microscopy) or for 24 h (for widefield epifluorescence microscopy).

### Comparative proteomics

For preparation of *in vitro* haptomonad attached flagellum sample, twenty-four Thermanox plastic coverslips on which *Leishmania* had attached were prepared in a 24 well plate. They were washed with 1 ml of Voorheis’s modified PBS (vPBS; 137 mM NaCl, 3 mM KCl, 16 mM Na_2_HPO_4_, 3 mM KH_2_PO_4_, 10 mM glucose, 46 mM sucrose, pH 7.6) three times and incubated in 1% (w/v) IGEPAL in PEME (0.1 M PIPES (pH 6.9), 2 mM EGTA, 1 mM MgSO_4_, 0.1 mM EDTA) with cOmplete, Mini Protease Inhibitor Cocktail without EDTA (Roche, Basel, Switzerland) for 5 min. Then coverslips were washed with 1 ml of a fresh 1% (w/v) IGEPAL in PEME with protease inhibitor cocktail and incubated in 1ml of 300 mM CaCl_2_ in PEME with protease inhibitor cocktail for 2 min. Coverslips were washed with PEME twice. For 24 coverslips, proteins were solubilised with 100 µl of a solubilisation buffer (2% (w/v) SDS, 60 mM Tris-HCl (pH 6.8), 50 mM DTT) with protease inhibitor cocktail per 6 coverslips and a total of ∼500 µl of attached flagellum sample was collected.

For promastigote non-attached flagellum sample preparation, 8 × 10^7^ promastigote cells were harvested by centrifugation (800 *g* for 7 min) and washed twice with 5 ml and 1 ml of vPBS. Cells were then incubated in 150 µl of 1% (w/v) IGEPAL in PEME with protease inhibitor cocktail and centrifuged (17,000 *g* for 2 min). The precipitate was resuspended in 150 µl of 300 mM CaCl_2_ in PEME with protease inhibitor cocktail and centrifuged (17,000 *g* for 10 min at 4°C). The precipitate was solubilised in 400 µl of a solubilisation buffer with protease inhibitor cocktail.

For mass spectrometry analysis, the haptomonad and promastigote samples were digested with sequencing grade trypsin (Promega, Southampton, UK) following, alkylation with iodoacetamide, immobilisation and clean up on suspension trap (S-Trap; ProTifi, Fairport, NY), following the manufacturer’s recommended protocol. Digestion proceeded overnight at 37°C before peptides were eluted, dried down and resuspended in aqueous 0.1% TFA for LC-MS/MS.

Peptides were loaded onto an mClass nanoflow UPLC system (Waters, Wilmslow, UK) equipped with a nanoEaze M/Z Symmetry 100 Å C_18_, 5 µm trap column (180 µm x 20 mm, Waters) and a PepMap, 2 µm, 100 Å, C_18_ EasyNano nanocapillary column (75 μm x 500 mm, Thermo Fisher Scientific, Waltham, MA). The trap wash solvent was aqueous 0.05% (v/v) trifluoroacetic acid and the trapping flow rate was 15 µL/min. The trap was washed for 5 min before switching flow to the capillary column. Separation used gradient elution of two solvents: solvent A, aqueous 0.1% (v/v) formic acid; solvent B, acetonitrile containing 0.1% (v/v) formic acid. The flow rate for the capillary column was 300 nL/min and the column temperature was 40°C. The linear multi-step gradient profile was: 3–10% B over 7 mins, 10– 35% B over 30 mins, 35–99% B over 5 mins and then proceeded to wash with 99% solvent B for 4 min. The column was returned to initial conditions and re-equilibrated for 15 min before subsequent injections.

The nanoLC system was interfaced with an Orbitrap Fusion Tribrid mass spectrometer (Thermo Fisher Scientific) with an EasyNano ionisation source (Thermo Fisher Scientific). Positive ESI-MS and MS^2^ spectra were acquired using Xcalibur software (version 4.0, Thermo Fisher Scientific). Instrument source settings were: ion spray voltage, 1,900 V; sweep gas, 0 Arb; ion transfer tube temperature; 275°C. MS^1^ spectra were acquired in the Orbitrap with: 120,000 resolution, scan range: *m/z* 375–1,500; AGC target, 4e^5^; max fill time, 100 ms. Data dependant acquisition was performed in top speed mode using a 1 s cycle, selecting the most intense precursors with charge states >1. Easy-IC was used for internal calibration. Dynamic exclusion was performed for 50 s post precursor selection and a minimum threshold for fragmentation was set at 5e^3^. MS^2^ spectra were acquired in the linear ion trap with: scan rate, turbo; quadrupole isolation, 1.6 *m/z*; activation type, HCD; activation energy: 32%; AGC target, 5e^3^; first mass, 110 *m/z*; max fill time, 100 ms. Acquisitions were arranged by Xcalibur to inject ions for all available parallelizable time.

Tandem mass spectra were extracted using MSConvert to .mgf format before submitting to database searching using Mascot (Matrix Science, version 2.7.0.1). Mascot was set up to search the *Leishmania mexicana* subset^21^ of the TriTrypDB database (https://tritrypdb.org/tritrypdb/app)^22^ appended with common proteomic contaminants (number of proteins in database = 8368). Data were searched with a fragment ion mass tolerance of 0.50 Da and a parent ion tolerance of 3.0 ppm. O-124 of pyrrolysine, j-16 of leucine/isoleucine indecision and carbamidomethyl of cysteine were specified in Mascot as fixed modifications. Oxidation of methionine was specified in Mascot as a variable modification. Scaffold (version Scaffold_5.2.0, Proteome Software Inc.) was used to validate MS/MS based peptide and protein identifications. Peptide identifications were accepted if they could be established at greater than 10.0% probability to achieve an FDR less than 1.0% by the Percolator posterior error probability calculation^23^. Protein identifications were accepted if they could be established at greater than 98.0% probability to achieve an FDR less than 1.0% and contained at least 2 identified peptides. Protein probabilities were assigned by the Protein Prophet algorithm^24^. Proteins that contained similar peptides and could not be differentiated based on MS/MS analysis alone were grouped to satisfy the principles of parsimony. Proteins sharing significant peptide evidence were grouped into clusters.

Among the proteins identified in the haptomonad and promastigote samples, those that were specific or more abundant in the haptomonad sample were listed. From the list, proteins that had known functions or localisations that would unlikely be involved in haptomonad adhesion were excluded e.g. mitochondrial. We also used TrypTag (http://tryptag.org/)^13^, the genome-wide protein localisation resource for the related parasite *Trypanosoma brucei* to remove proteins whose orthologs localised to organelles unlikely to be involved in adhesion. The top 20 proteins in the list thus selected were then endogenously tagged with mNeonGreen and their localisation in a promastigote and haptomonad was examined.

### Scanning electron microscopy

Intact *in vitro* haptomonad, or haptomonad after being treated with 1% IGEPAL in PEME or with 1% IGEPAL and 300 mM CaCl_2_ in PEME on Thermanox plastic coverslips were fixed respectively with 2.5% glutaraldehyde in PEME. After an hour, coverslips were washed once in PEME and once in ddH_2_O. The coverslips were then dehydrated using increasing concentrations of ethanol (30%, 50%, 70%, 90%, 100% (v/v), and 2 × absolute ethanol; 10 min / step)). The coverslips were then critical point dried, mounted onto SEM stubs using carbon stickers, and sputter coated with a layer of 12–14 nm of gold. Images were taken on a Hitachi S-3400N scanning electron microscope at 5 kV, at a 5.5 mm working distance.

### Transfection and drug selection

Generation of *L. mexicana* tagging constructs and sgRNA templates for endogenous mNeonGreen or mCherry tagging were generated by the PCR method as previously described using pLPOT (mNG/Puromycin) or pLPOT (mCh/Neomycin) as the template, respectively^25^. Transfection of cells was performed as previously described using the Amaxa Nucleofector-2b^26^. Primers for constructs and sgRNA were designed using LeishGEdit (http://www.leishGEdit.net)^27^. For the generation of KIAP1, 2 and 3 null mutants using CRISPR/Cas9 mediated genome editing, the C9/T7 cell line was transfected with guide and repair constructs generated by PCR using primers designed on the LeishGEdit using the G00 primer and the pTBlast and pTPuro plasmids as templates^27^. Constructs were transfected using a Nucleofector 2b. Successful transfectants were selected with 20 µg/ml Puromycin (Melford Laboratories, Ipswich, UK) or 20 µg/ml G-418 (Melford Laboratories), or 5 µg/ml Blasticidin (Melford Laboratories) and 20 µg/ml G-418.

### Generation of KIAP1–3 add back cell lines

To produce the add back cell lines, the *KIAP1* (*LmxM.01.0010*), *KIAP2* (*LmxM.30.0390*) and *KIAP3* (*LmxM.20.1190*) genes were PCR amplified and then cloned into the *HindIII* and *SpeI* or *SpeI* and *BamHI* restriction sites of the constitutive expression plasmid (pJ1364) to generate a C-terminally tagged version of KIAP1 and KIAP2 with an mNG and a triple myc tag or an N-terminally tagged version of KIAP3 with a triple myc tag, respectively, as described previously^28,29^. The plasmids were linearised by digestion with PacI (New England Biolabs, Ipswich, UK) and ethanol precipitated before transfection. The constructs were transfected using a Nucleofector 2b. Successful transfectants were selected with 25 µg/ml phleomycin (Melford Laboratories).

### Light microscopy

For light microscopy of living cells, haptomonad cells attached to a piece of a gridded glass coverslip were washed twice in DMEM, incubated in DMEM with Hoechst 33342 (1 µg/ml) for 5 min and then washed twice in DMEM. Coverslip pieces were mounted onto another glass coverslip and then onto a glass slide, with the cell attachment side facing up. Attached cells were imaged using a Zeiss ImagerZ2 microscope (Carl Zeiss, Jena, Germany) with Plan-Apochromat 63×/1.4NA Oil objective and a Hamamatsu Flash 4 camera (Hamamatsu Photonics, Hamamatsu, Japan).

### Bioinformatic analysis

The 18S rRNA sequences of the selected kinetoplastids were retrieved from TriTrypDB. Alignment and phylogenetic reconstructions of the 18S rRNA sequences of the kinetoplastids were performed using the function “build” of ETE3 3.1.2^30^ as implemented on the GenomeNet (https://www.genome.jp/tools/ete/). Alignment was performed with MAFFT v6.861b with the default options^31^. Columns with more than one percent of gaps were removed from the alignment using trimAl v1.4.rev6^32^. ML tree was inferred using RAxML v8.2.11 ran with model GTRGAMA and default parameters^33^. Branch supports were computed out of 100 bootstrapped trees.

Reciprocal best BLAST was used to find orthologous sequences^34^. *L. mexicana* KIAP protein sequences were used to interrogate the protein sequence database on TriTrypDB by BLASTP, which identified sequences in other kinetoplastid species. The top hit from each species was then used to interrogate the *L. mexicana* genome by BLASTP and if this returned the same *L. mexicana* KIAP protein as used in the initial search the two proteins were considered orthologous. KIAP sequences were analysed by the InterPro (https://www.ebi.ac.uk/interpro/). Protein structures were predicted using AlphaFold^35–37^, using the exact pipeline previously described^37^. Visualisation and superposition of protein structures were carried out with PyMOL (DeLano Scientific LLC, San Carlos, CA, http://www.pymol.org). The structure of human Tetranectin was obtained from the RCSB protein data bank (https://www.rcsb.org/; PDB: 1HTN)^38^.

### Time-lapse observation

For time-lapse observation, log phase promastigotes (1 × 10^6^ cells/ml) were cultured in a µ- dish 35 mm, high glass bottom (iBidi, Gräfelfing, Germany) for 12 h at 28°C with 5% CO_2_, and the dish was washed five times with fresh M199 medium before starting imaging. Cells about to adhere to the glass were recorded using Zeiss LSM 880 confocal microscope (Carl Zeiss, Jena, Germany) with Plan-Apochromat 63×/1.4NA Oil objective and 488 and 561 nm lasers (1% laser powers) at 28°C with 5% CO_2_ in a chamber with controlled temperature and CO_2_ concentration.

### Confocal microscopy

Log phase promastigotes (1 × 10^6^ cells/ml) were cultured in a µ-dish 35 mm, high glass bottom for 24 h, and the dish was washed more than five times with fresh M199 medium to remove as many non-adherent cells as possible before starting imaging. High resolution confocal microscopy images were acquired with a Zeiss LSM880 with Airyscan using a Plan-Apochromat 100×/1.46NA Oil DIC objective, with 488 and 561 nm lasers. Confocal z-stacks were acquired in superresolution mode using line scanning and the following settings: 40 × 40 nm pixel size, 50 nm z-step, 0.66 µs/pixel dwell time, 800 gain, 2% laser power. The z-stack images were processed and analysed using Zen Black software (Carl Zeiss, Jena, Germany) and Fiji^39^.

### Analysis of serial section electron microscopy tomography data sets

The tomography data sets that we deposited on the Electron Microscopy Image Archive (EMPIAR; https://www.ebi.ac.uk/empiar/)^40^ in our previous study (EMPIAR-11467 and EMPIAR-11468)^11^ were re-used for the 3D structural analysis of the attachments of the *in vitro* and *in vivo* haptomonad cells. Three-dimensional models of the electron tomography data were created using 3dmod (IMOD software package)^41^ as previously described^11^.

### Fluorescence recovery after photobleaching

Log phase promastigotes (KIAP1::mNG, SMP1::mCh, 1 × 10^6^ cells/ml) were cultured in a µ-dish 35 mm, high glass bottom for 24 h, and the dish was washed more than five times with fresh M199 medium to remove as many non-adherent cells as possible before starting imaging. A Zeiss LSM880 with Airyscan was used for imaging and simultaneous photobleaching. The pre-bleach image of the attached flagellum was acquired using a Plan-Apochromat 100×/1.46NA Oil DIC objective, with 488 and 561 nm lasers. A ROI including part of KIAP1::mNG or SMP1::mCh fluorescence signal on a glass surface was imaged 5 times (short-time observation) or twice (long-time observation) before bleaching. KIAP1::mNG or SMP1::mCh was bleached using the 488 or 561 nm laser, respectively (two or ten iterations of full laser power for the 488 or 561 nm laser, respectively, fluorescence reduction of ∼60%). Fluorescence recovery in the KIAP1::mNG or SMP1::mCh was assessed by imaging once every 0.24 s for up to 9.84 s (short-time observation) or every 20 s for up to 300 s (long-time observation) using the 488 and 561 nm low power laser excitation (0.5%). Image analysis was performed using Zen Black software and Fiji. Normalised fluorescence values were calculated by dividing fluorescence value at each time point by average fluorescence value before bleaching and plotted with Microsoft Excel.

### *In vitro* attachment quantification

Log phase promastigotes (5 × 10^6^ cells/ml) of parental (SMP1::mCh), KIAP1, 2 and 3 knockout and add back cell lines were cultured on ∼5 × 5 mm pieces of gridded glass coverslips grid-500 (iBidi) in a 24 well plate with 1 ml of complete M199 medium for 24 h at 28°C with 5% CO_2._ The coverslips were washed twice with 1 ml of DMEM, incubated in 1 ml of DMEM with Hoechst 33342 (1 µg/ml) for 5 min, and washed twice with 1 ml of DMEM. The glass pieces were mounted with another glass coverslip on a glass slide. Cells attached in one grid area of 500 µm × 500 µm were imaged using a Zeiss ImagerZ2 microscope with Plan-Apochromat 20×/0.8NA objective and Hamamatsu Flash 4 camera. The number of cells attached in one grid area was measured based on the phase or bright field image and Hoechst signal using Fiji.

### Sand fly infection experiments

Females of *Lutzomyia longipalpis* were fed through a chick-skin membrane on heat-inactivated sheep blood containing *Leishmania mexicana* promastigotes (Parental (SMP1::mCh), KIAP1 and 3 knockout and add back cell lines) from log-phase cultures (day 3–4) at a concentration 1 × 10^6^ cells/ml. Blood-engorged females were separated and maintained at 26°C with free access to 50% sugar solution. On days 6 and 9 post-blood meal (PBM) females were dissected in drops of saline solution. The individual guts were checked for presence and localisation of *Leishmania* under the light and fluorescence microscope Olympus BX51. Special emphasis was given to the colonisation of the stomodeal valve. Levels of *Leishmania* infections were graded into four categories: negative, light (<100 parasites/gut), moderate (100–1000 parasites/gut) and heavy (>1000 parasites/gut). The experiment was repeated twice.

### Immunofluorescence microscopy

Log phase promastigotes (1 × 10^6^ cells/ml) of KIAP3 add back cell line which expresses 3myc::KIAP3 were cultured on 18 × 18 mm precision cover glasses thickness No. 1.5H (Paul Marienfeld, Lauda-Königshofen, Germany) which were sterilised with 100% ethanol in a 6 well plate with 3 ml of complete M199 medium for 24 h at 28°C with 5% CO_2._ The coverslips were washed with 3 ml of DMEM, treated with 1% IGEPAL in PEME for 1 min and fixed for 30 min with 4% (w/v) paraformaldehyde in PEME, washed with phosphate-buffered solution (PBS) three times for 5 min, blocked for 30 min with 1% BSA in PBS at RT. The coverslips were then incubated with c-Myc monoclonal antibody (9E10; Invitrogen, Paisley, UK) in PBS containing 1% BSA (1:100 dilution) overnight at 4°C, washed with PBS three times for 5 min, incubated with Alexa Fluor 546-conjugated goat anti-mouse secondary antibody (Invitrogen) in PBS containing 1% BSA (1:1000 dilution) for 1 h at RT and washed with PBS three times for 5 min. Then, the coverslips were treated with 0.1 µg/ml Hoechst 33342 in PBS for 5 min, washed with PBS twice and mounted with VECTASHIELD mounting medium for imaging using a Zeiss Imager Z2 microscope with the Plan-Apochromat 63×/1.4NA Oil objective and Hamamatsu Flash 4 camera.

## Statistical analysis

Means, SDs and SEMs were calculated using Microsoft Excel. Statistical significance was determined using two-tailed Welch’s *t-test* carried out with Microsoft Excel. Differences were considered significant at the level of p<0.05. Data were plotted with Microsoft Excel or the Matplotlib package in Python^42^.

## Data availability

The data and statistics that support the findings of this study are available as Source Data files.

## Acknowledgements

LC-MS/MS was performed using equipment at the University of York Centre of Excellence in Mass Spectrometry, created thanks to a major capital investment through Science City York, supported by Yorkshire Forward with funds from the Northern Way Initiative, and subsequent support from the Engineering and Physical Sciences Research Council (EP/K039660/1; EP/M028127/1). We thank Dr Richard Wheeler (University of Oxford) for the support in the analysis using the AlphaFold and providing the sand fly cartoon in Fig. 4b; the Oxford Brookes Centre for Bioimaging for the support in light and electron microscopy; Prof Keith Gull (University of Oxford), Prof Sue Vaughan (Oxford Brookes University) and members of the Sunter and Vaughan laboratory for discussions and comments on the manuscript. RY was supported by a JSPS Overseas Research Fellowship and NIBB Collaborative Research Program (20-515). This project has also received resources funded by the European Union’s Horizon 2020 research and innovation programme under grant agreement No 731060 (Infravec2). Work in the lab of JDS is supported by the Wellcome Trust (221944/Z/20/Z). JS, BV, KP and PV were supported by ERD funds, project CeRaViP (16_019/0000759) and Czech Science Foundation (GACR 21-15700S).

## Author contributions

Conceptualisation, RY, JDS; Formal Analysis, RY, KP, Funding, RY, PV, JDS; Investigation, RY, KP, ER, FML, AT, SN, JS, BV; Supervision, SN, PV, JDS; Visualisation, RY, KP; Writing, RY, KP, JS, PV, JDS.

## Figures

**Extended Data Fig. 1.**
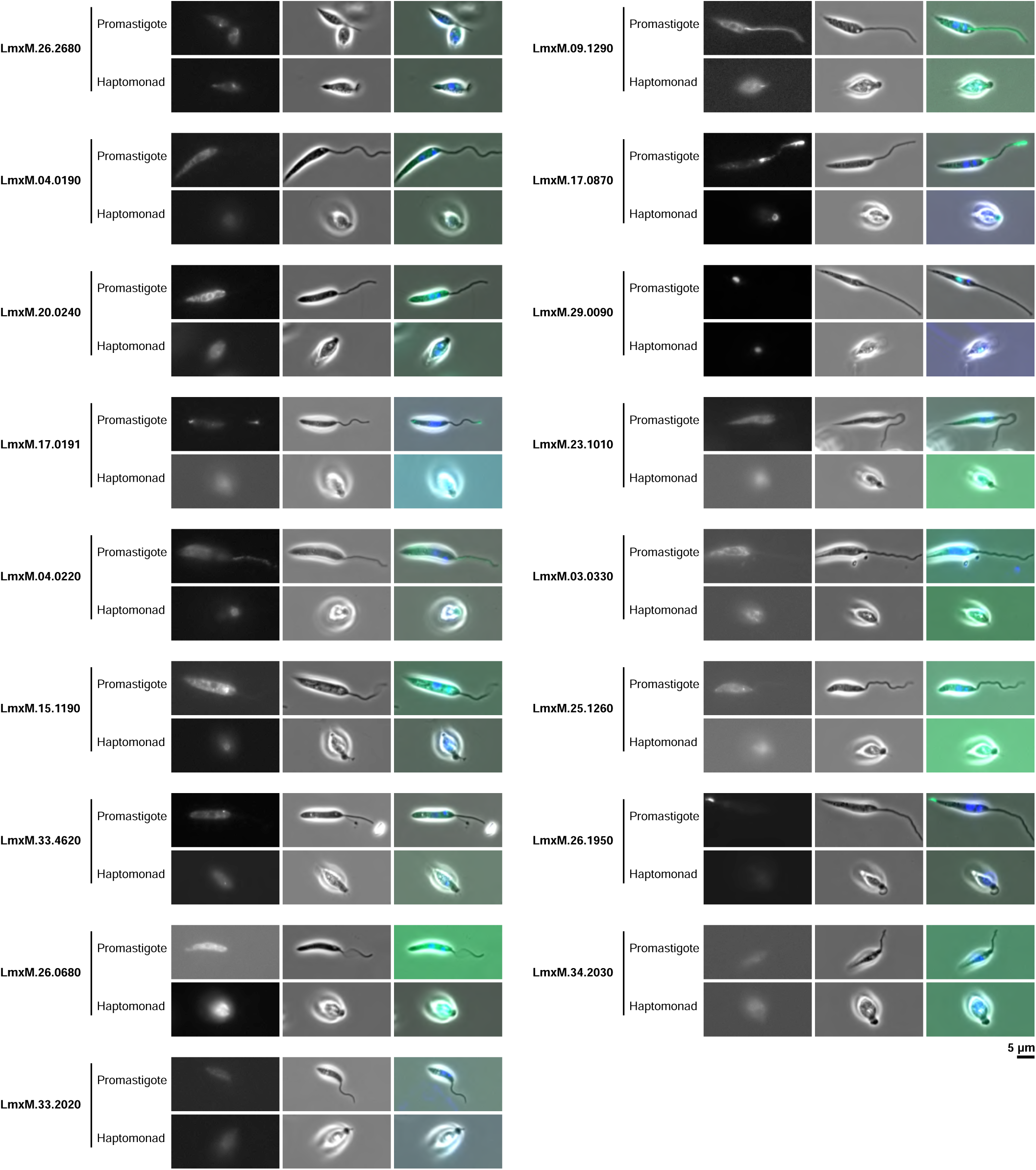
mNeonGreen tagging screening of the identified proteins in the comparative proteomics analysis. Localisation of the identified proteins in the comparative proteomics which are endogenously tagged with mNG in promastigote and *in vitro* haptomonad are shown.

**Extended Data Fig. 2.**
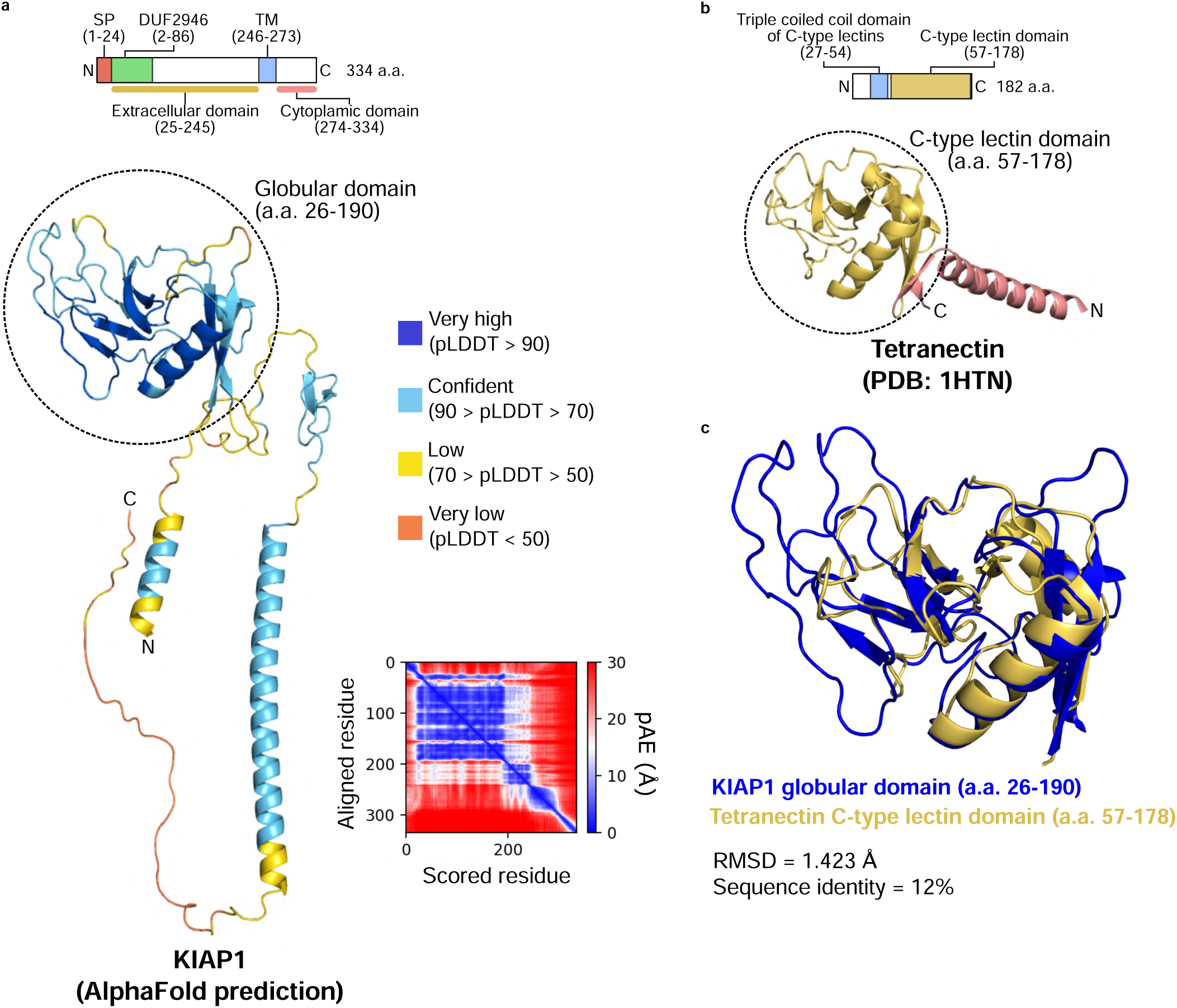
AlphaFold prediction shows a globular domain of KIAP1 has a similar 3D structure with that of a C-type lectin domain. **a**, Domain structure and 3D structure of KIAP1 predicted by AlphaFold. Globular domain of KIAP1 is indicated by a dotted circle (amino acids (a.a.) 26-190). Colours of the 3D structure represent the predicted local distance difference test score (pLDDT) of the AlphaFold prediction. The corresponding plot of predicted aligned error (pAE) of the residues is shown at bottom right. **b**, Domain structure and 3D structure of human Tetranectin (PDB: 1HTN). C-type lectin domain is shown in yellow in the domain and 3D structure. **c**, Superimposition of the 3D structures of the KIAP1 globular domain (blue; a.a. 26–190) and Tetranectin C-type lectin domain (yellow; a.a. 57– 178). The root-mean-square deviation (RMSD) between two protein structures calculated using PyMOL was 1.423 Å and the sequence identity was 12%.

**Extended Data Fig. 3.**
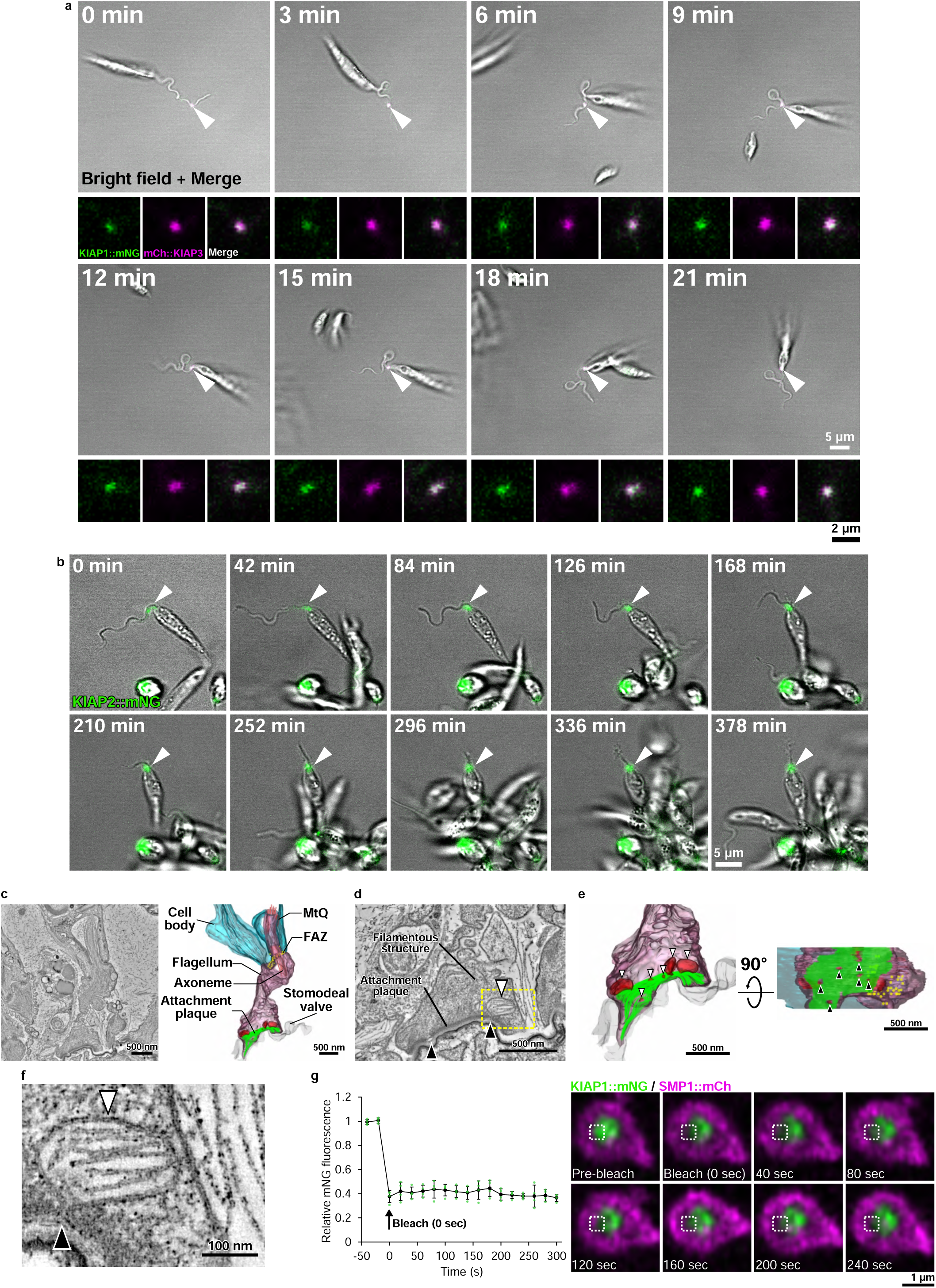
Time-lapse observation of the development of KIAP1, 2 and 3 during *in vitro* haptomonad adhesion processes. **a**, Sequential frames (at 3 min intervals) from a time-lapse video of adhesion of a haptomonad cell and development of KIAP1::mNG and mCh::KIAP3. The images above show merged images of a bright field image and mNG and mCh signals. Note that the starting point of adhesion (arrowheads) remains fixed during the adhesion process. The images presented below at each time point represent the confocal microscopy images of KIAP1::mNG (green) and mCh::KIAP3 (magenta), respectively, as well as their merged images. **b**, Sequential frames (at ∼40 min intervals) from a time-lapse video of a complete adhesion process of a haptomonad cell and development of KIAP2::mNG. Note that the starting point of adhesion (arrowheads) remain fixed until the completion of adhesion. **c**, A slice of a serial tomogram and 3D reconstruction of an *in vivo* haptomonad cell in the sand fly. Vesicles near the attachment plaque (green) are shown in red. The name of each structure is given in the 3D reconstruction. **d**, Magnified view of the tomogram showing that multiple vesicles (white arrowheads) are seen near the attachment plaque and the plaque is interrupted at the fusion of the vesicle and flagellar membrane (black arrowheads). **e**, Magnified view of the 3D reconstruction showing that multiple vesicles (white arrowheads) are seen near the attachment plaque (green) and the plaque is interrupted at the fusion of the vesicle and flagellar membrane (black arrowheads). Side (left) and bottom (right) views of the attached flagellum are shown. **f**, Magnified view of the vesicle near the attachment plaque (yellow dotted box in d). Note that structures resembling a folded lipid bilayer or a filamentous extracellular matrix around the haptomonad in the sand fly are seen. **g**, Long-time FRAP experiment of KIAP1::mNG. The relative fluorescence intensity changes before and after photobleaching is shown in the line graph, with the average fluorescence intensity before photobleaching as 1. The timing of photobleaching is indicated by an arrow in the graph. Data represent mean ± SD (n = 4 independent experiments). Values from each experiment are shown with green dots, respectively. Sequential frames (at 40 sec intervals) of confocal microscopy observation of KIAP1::mNG and SMP1::mCh are shown. The area of photobleaching is indicated with dotted white boxes.

**Extended Data Fig. 4.**
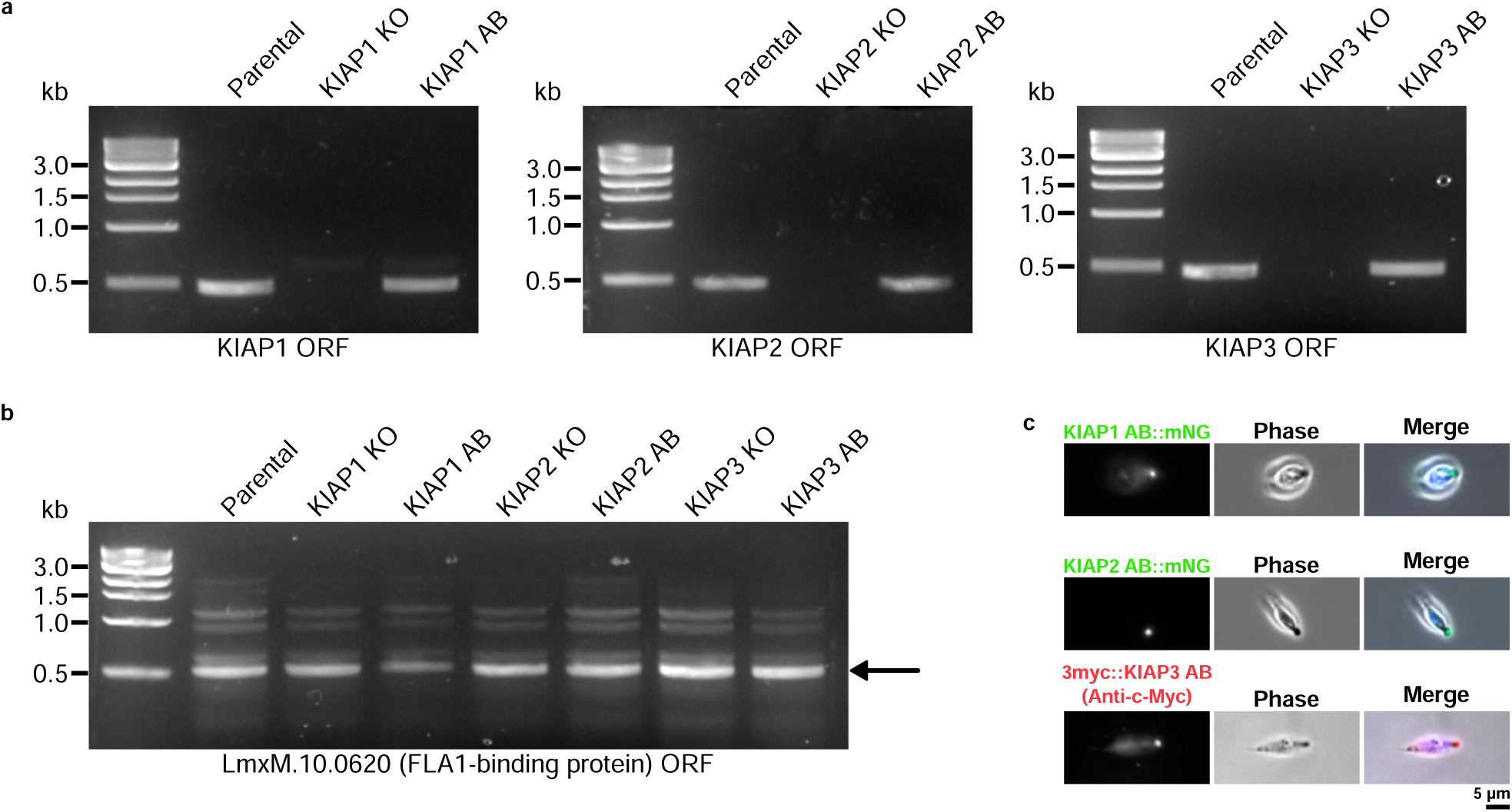
Confirmation of knockout and add back of KIAP1, 2 and 3 genes. **a**, gDNA from parental and KIAP1, 2 and 3 knockout and add back cells was analysed by PCR. PCR confirmed that KIAP1, 2 and 3 open reading frame (ORF) was no longer present in knockout cells, and KIAP1, 2 and 3 ORF was present in add back cells. **b**, The presence of the ORF of FLA1-binding protein (LmxM. 10.0620) was confirmed by PCR using the respective gDNA. The black arrow indicates the amplified 500 bp bands from the ORF of FLA1-binding protein. **c**, Fluorescence microscopy confirmed that the KIAP1::mNG::3myc, KIAP2::mNG::3myc and 3myc::KIAP3 expressed in add back cell localised to the attached flagellum of *in vitro* haptomonds. Note that to mitigate the reduced attachment capacity of *L. mexicana* cell line expressing KIAP3 N-terminally tagged with mNG and a triple myc tag, KIAP3 N-terminally tagged only with a triple myc tag was re-introduced in KIAP3 AB cells. Localisation of 3myc::KIAP3 in a haptomonad cell was confirmed by immunofluorescence microscopy using c-Myc monoclonal antibody (9E10) and Alexa Fluor 546-conjugated goat anti-mouse secondary antibody.

**Supplementary Dataset 1. Mass spectrometry data of the haptomonad and promastigote samples.**

**Supplementary Dataset 2. List of proteins selected for the localisation screening with mNeonGreen tagging.**

**Supplementary Video 1. Time-lapse video of a complete adhesion process of an *in vitro* haptomonad cell expressing mNG::KIAP3.** Playback of ∼3 h at 500x speed.

**Supplementary Video 2. Time-lapse video of an initial adhesion process of an *in vitro* haptomonad cell expressing KIAP1::mNG and mCh::KIAP3.** Playback of ∼25 min at 100x speed.

**Supplementary Video 3. Time-lapse video of an initial adhesion process of an *in vitro* haptomonad cell expressing KIAP2::mNG.** Playback of ∼15 min at 100x speed.

**Supplementary Video 4. Time-lapse video of a complete adhesion process of an *in vitro* haptomonad cell expressing KIAP2::mNG.** Playback of ∼6 h at 1200x speed.

